# Validation of a heat-inducible *Ixodes scapularis* HSP70 promoter and developing a tick-specific 3xP3 promoter sequence in ISE6 cells

**DOI:** 10.1101/2023.11.29.569248

**Authors:** Michael Pham, Hans-Heinrich Hoffmann, Timothy J. Kurtti, Randeep Chana, Omar Garcia-Cruz, Simindokht Aliabadi, Monika Gulia-Nuss

## Abstract

*Ixodes scapularis* is an important vector of many pathogens, including the causative agent of Lyme disease, tick-borne encephalitis, and anaplasmosis. The study of gene function in *I. scapularis* and other ticks has been hampered by the lack of genetic tools, such as an inducible promoter to permit temporal control over transgenes encoding protein or double-stranded RNA expression. Studies of vector-pathogen relationships would also benefit from the capability to activate anti-pathogen genes at different times during pathogen infection and dissemination. We have characterized an intergenic sequence upstream of the heat shock protein 70 (HSP70) gene that can drive *Renilla* luciferase expression and mCherry fluorescence in the *I. scapularis* cell line ISE6. In another construct, we replaced the *Drosophila melanogaster* minimal HSP70 promoter in the synthetic 3xP3 promoter with a minimal portion of the *I. scapularis* HSP70 promoter and generated an *I. scapularis* specific 3xP3 (Is3xP3) promoter. Both promoter constructs, IsHSP70 and Is3xP3, allow for heat-inducible expression of mCherry fluorescence in ISE6 cells with an approximately 10-fold increase in the percentage of fluorescent positive cells upon exposure to a 2 h heat shock. These promoters described here will be valuable tools for gene function studies and temporal control of gene expression, including anti-pathogen genes.

**Graphical abstract:** 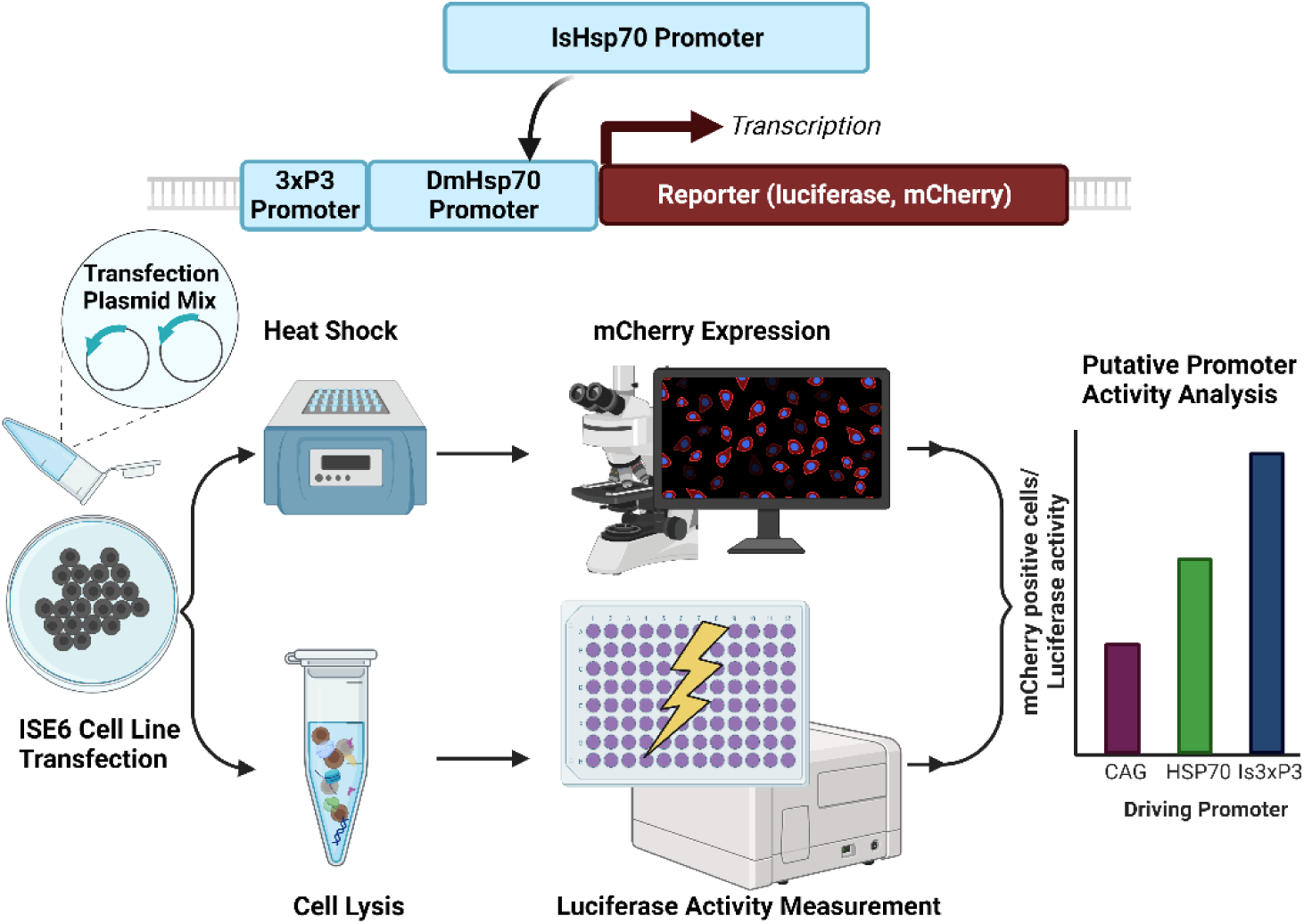

## INTRODUCTION

Ticks are obligate hematophagous parasites and are important vectors of a wide variety of pathogens (***Gulia-Nuss et al., 2016***). Lyme disease (LD), caused by the spirochete *Borrelia burgdorferi* and vectored by the black-legged tick, *Ixodes scapularis,* is the most prevalent vector-borne disease in the United States. The Centers for Disease Control and Prevention (CDC) estimates 476,000 cases of LD every year (***Centers for Disease Control and Prevention, National Center for Emerging and Zoonotic Infectious Diseases (NCEZID), Division of Vector-Borne Diseases, December 21, 2018***). Despite their importance, our knowledge of the biology of ticks on a molecular level is limited. Advances in tick genomics and genetics have mainly been stymied by a lack of molecular tools for forward genetics. This is in contrast to insects for which numerous transgenic development and gene editing tools are available. CRISPR/Cas9 (clustered, regularly interspaced, short palindromic repeat/CRISPR-associated protein 9) is revolutionizing genome editing in non-model organisms. We have made significant progress in optimizing tick embryo injections and developing CRISPR gene knockout strategies in ticks (***Sharma et al., 2022***); however, knock-in through homologous dependent repair (HDR) depends on developing reporter constructs using specific promoters. Promoters from *Drosophila melanogaster* genes such as polyubiquitin (***Lee et al., 1988***), Actin5C (***Stebbins et al., 2001***), HSP70 (***Lis et al., 1983***), and the artificial 3xP3 promoter (***Berghammer et al., 1999***) have been successfully used to express marker genes through transposase or CRISPR-mediated knock-in in a wide variety of insects and other organisms (***Atkinson et al., 2001; Berghammer et al., 1999; González-Estévez et al., 2003***). While these exogenous promoters have proven to be widely applicable, they are not universal, and our previous experiments showed that they were not functional in ticks.

Because of the unavailability of promoters and the lack of application of exogenous promoters, there is an urgent need to identify endogenous and non-endogenous promoters that may function across multiple tick species. Several tick endogenous promoters such as *Haemophysalis longicornis* Ferritin (HlFerritin), HlActin, *I. scapularis* ribosomal protein L4 (Isrpl4), *Rhiphicephalus microplus* rpl4(Rm-rlp4), Rm-EF-1α, Is-microsomal glutathione S-transferase, Is-ribosomal protein S24, and RmPyrethroid metabolizing esterase gene (***Hernandez et al., 2019; Kusakisako et al., 2018; Machado-Ferreira et al., 2015; Naranjo et al., 2013; Shi et al., 2022; Tuckow and Temeyer, 2015***) as well as non-endogenous promoters (human phosphoglycerate kinase, CAG, cytomegalovirus [CMV], polyhedrin promoter, Cauliflower Mosaic Virus) (***Esteves et al., 2008; Kurtti et al., 2008; Machado-Ferreira et al., 2015; Naranjo et al., 2013a; Shi et al., 2022; Tuckow and Temeyer, 2015; You et al., 2003***) have been demonstrated to be functional in tick cell lines; however only one (HlFerritin) is inducible (with ferrous sulfate). None of the endogenous tick promoters are available to other researchers. Identifying effective and consistent inducible promoter or enhancer sequences for use in driving transient transgene expression would broaden the types of experiments that could be performed using tick cell lines and permitting temporal control of gene expression. Therefore, we aimed to identify tick-specific promoters that could be beneficial for gene expression, RNAi, or knock-in studies. Here, we demonstrate functional and heat-inducible *I. scapularis* heat shock protein 70 (IsHSP70) promoter and an artificial *I. scapularis* specific 3xP3 promoter under the control of a minimal IsHSP70 core promoter (Is3xP3).

## METHODS

We synthesized a 1,301 bp sequence fragment (GeneUniversal) upstream of an *I. scapularis* HSP70 (ISCW024910/ISCGN274230) (Figure 1) including the entire 5’ untranslated region of the gene (UTR) with restriction enzyme sites added to the 5’ and 3’ end (HSP70, Acc65I-SacI, and BglII), PCR (primers listed on Table 1) amplified with Taq polymerase PCR and cloned them into the pGEM-T-easy vector. Putative promoter sequences were digested and ligated into pGL-4.79 [Rluc] (Promega), a *Renilla* luciferase reporter construct lacking a promoter.

**Figure 1:**
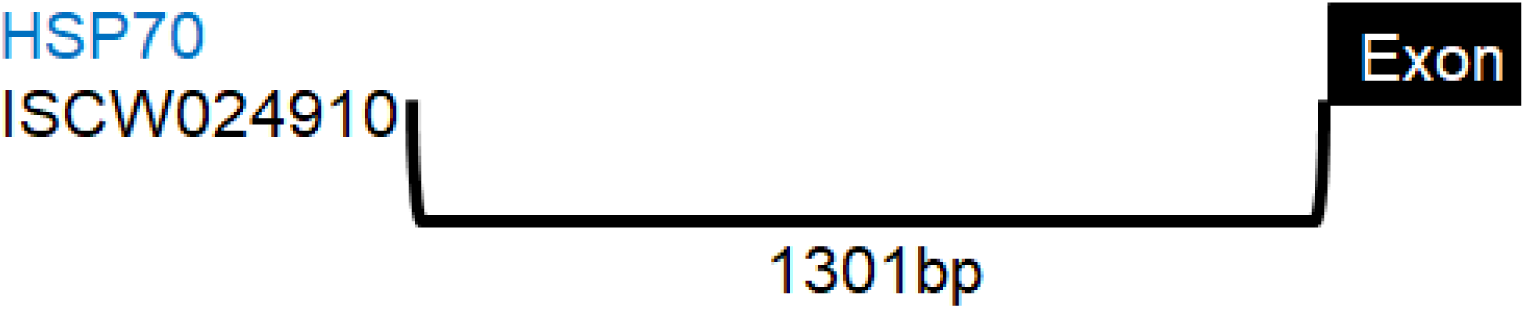
A gene map of the HSP70 (ISCW024910/ ISCGN274230 promoter sequence. This gene only has one exon. Upstream region including the entire 5’ UTR was cloned into the pGL-4.79 [Rluc] vector.

**Table 1:**
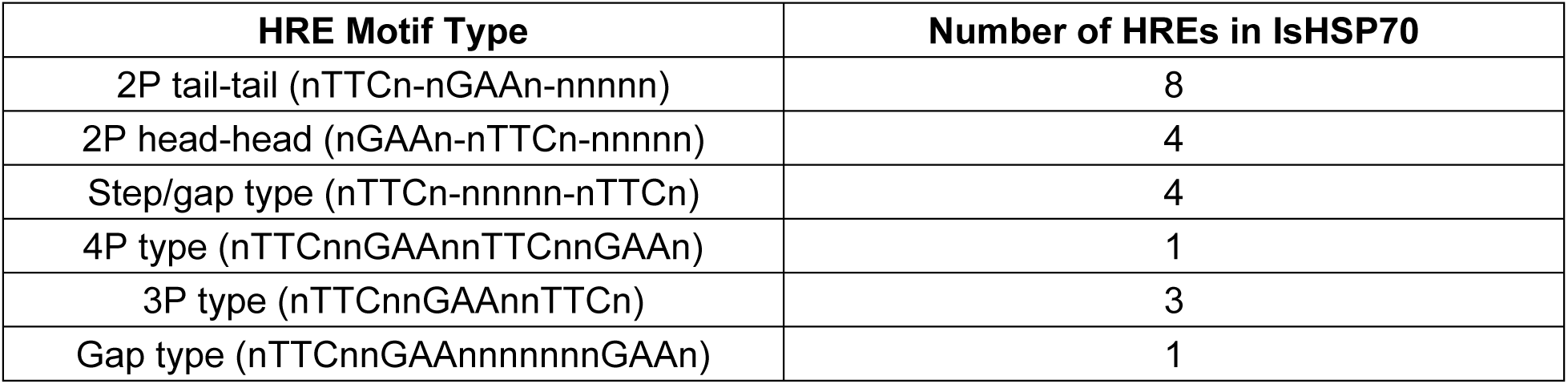
Heat Response Elements (HRE) motif types and number in *Ixodes scapularis* HSP70 gene.

*Renilla* luciferase was swapped from pGL-4.79 [Rluc] and its associated 3’ SV40 polyadenlyation sequence with EGFP (with its SV40 polyadenlyation sequence) through restriction enzyme cloning (gift from Dr. Shengzhang Dong, Johns Hopkins University).

mCherry was subcloned from a mCherry plant expression plasmid (gift from Dr. Jeffrey Harper, University of Nevada, Reno), using primers containing a BglII site and Kozak sequence for the 5’ primer and a MfeI site for the 3’ primer. The restriction sites were used to insert mCherry and remove EGFP from the promoterless pGL4.79-EGFP construct we generated earlier. We retained the SV40 sequence from EGFP (GeneUniversal). pGL-4.79 vectors contain double SfiI sites that flank the multiple cloning sites (the site of promoter insertion). HSP70 sequences were subcloned from their current Rluc constructs into pGL-4.79 mCherry (GeneUniversal).

The sequences of 3xP3 with the three Pax6 binding regions and the *Drosophila melanogaster* HSP70 minimal promoter (***Berghammer et al., 1999; Thurmond et al., 2019***)) were determined. The *Drosophila* minimal promoter (TATA box to start codon, 123 bp) from 3xP3 (***Berghammer et al., 1999***) was replaced with the minimal promoter from our IsHSP70 construct (TATA box to start codon, 453 bp) (GeneUniversal).

### Cell lines

ISE6 cells (*Ixodes scapularis*; sex: unspecified; embryonic origin) were obtained from Prof. Ulrike G. Munderloh (University of Minnesota, MN) and cultured in Leibovitz’s L-15C-300 medium (pH = 7.25) supplemented with 5% fetal bovine serum (FBS), 5% tryptose phosphate broth (TPB) and 0.1% bovine lipoprotein cholesterol (BLC) concentrate at ambient atmosphere and 32 °C (***Munderloh et al., 1994; Munderloh and Kurtti, 1989***). Cells were tested negative for contamination with mycoplasma.

### Transfection

#### ISE6 cell plasmid transfection

ISE6 cells were seeded into 96-well plates at 0.1 mL/well of 5×10^5^ cells/mL and incubated at 32 °C overnight. The following day, plasmid DNA was diluted into 10 μL Opti-MEM and mixed with 10 μL Opti-MEM containing Lipofectamine™ MessengerMAX™ transfection reagent (ThermoFisher Scientific: LMRNA001) at a 1:1 ratio of plasmid DNA [μg] to transfection reagent [μL]. The transfection mix (20 μL) was incubated for 15 min at RT and added to wells containing 30 μL medium, followed by spin-transfection for 1 h at 1,000 *g* at 32 °C. After 6 h, the transfection mix was replaced with 100 μL fresh L-15C-300 medium. Cells were incubated for 48 h before being exposed to a 2 h heat shock at 37 °C or 40 °C. At 72 h post-transfection (24 h post heat shock), cells were fixed by adding 100 μL of 7% formaldehyde to each well. Nuclei were stained with Hoechst 33342 (ThermoFisher Scientific: 62249) at 1 µg/mL. Images were acquired with a fluorescence microscope and analyzed using ImageXpress Micro XLS (Molecular Devices, Sunnyvale, CA).

#### Luciferase activity

ISE6 cells were transfected with Renilla luciferase constructs using Lipofectamine 3000 (Thermo Fisher Scientific Inc, USA). ISE6 cells were seeded into 12.5 cm^2^ flasks at 2 × 10^6^ cells/mL and incubated at 32 °C overnight. Prior to transfection, cell layers were rinsed with Dulbecco’s PBS and held in 1 mL of serum-free L15Cd (SFL15Cd). Lipofectamine 3000 (7.5 µL) in 125 µL SFL15Cd was mixed with 125 µL SFL15Cd containing P3000 (10 µL) and 5 µg of plasmid DNA. Plasmid, in 250 µL complete Lipofectamine 3000 mixture, was incubated at room temperature for 20 min and then added dropwise onto cell layers. Cell layers were gently rocked for 1 h at 32 °C, and 2 mL of complete growth medium was added. Transfected cultures were incubated at 32 °C for 7-9 days and evaluated for luciferase activity. Cell layers were washed with Dulbecco’s PBS and incubated in 500 µL luciferase assay lysis buffer for 15 min. Lysates were stored at −20 °C until evaluated. Lysates were examined for luciferase activity using a commercial kit assay (Promega Cat.# E2810). Lysates (20 µL) were inoculated into wells of a 96 well plate, and Renilla Luciferase Assay Reagent (100 µL/well) was added. Fluorescence was measured at 10 seconds at Relative Luciferase Units (RLU) using a luminometer microplate reader (Biotek synergy H1 hybrid plate reader with Gen5v. Software, Biotek, Winooski, VT, USA).

## RESULTS

### HSP70 Promoter

Following a previously published strategy (***Anderson et al., 2010; Carpenetti et al., 2012; Gross et al., 2009***) we identified the HSP70 genes in the fruit fly, *Drosophila melanogaster,* and mosquito, *Aedes aegypti*, that are ubiquitously expressed across tissues and life stages (Uniprot.org). Using blastp, we identified a putative ortholog of HSP70: ISCW024910 in the *I. scapularis Wikel* genome with greater than 70% amino acid sequence identity and consisted of a single exon. We confirmed this sequence with our highly contiguous genome assembly (***Nuss et al., 2023***). We located an intergenic sequence 5’ from the coding region of these genes and confirmed potential promoter function by bioinformatics analysis (data not shown) (***Shahmuradov and Solovyev, 2015; Solovyev and Shahmuradov, 2003***) (Figures 1, 2). We also confirmed the heat response element sequences (IsHSP70) and arthropod initiation factor motifs (***Cherbas and Cherbas, 1993***) (Table 1 and Figure 2).

**Figure 2:**
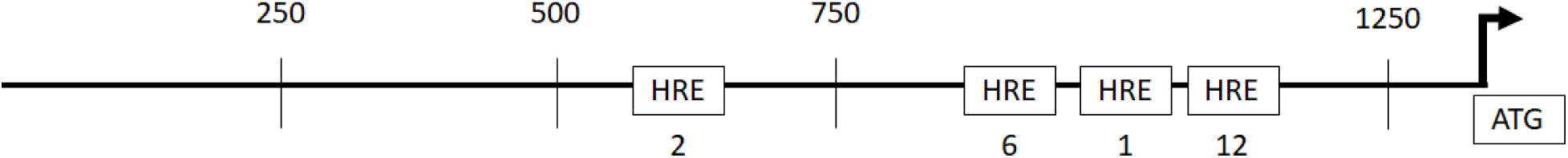
A gene map of the cloned putative promoter region of HSP70 (1,315 base pairs in size) with heat response element locations marked. Numbers below HRE locations indicate the numbers present at that location. (HRE: Heat Response Element)

The sequence upstream of the predicted IsHSP70 gene was searched to identify putative heat regulatory elements (HREs) sequences (sequence motifs listed in Table 1) (***Carpenetti et al., 2012; Sorger, 1991***). HREs are regions where heat shock factors (HSF) bind and activate HSP70 transcription and regulate the HSP70 gene, and thus, an organism’s response to heat stress is directly affected by HREs. These HRE sequence motifs are conserved across many species and if clustered, can act cooperatively (***Chatterjee and Burns, 2017; Sakurai and Enoki, 2010***). Based on the conserved HRE sequence motifs, we identified 21 potential HREs in a single general area 200 - 700 bp 5’ from the start of the HSP70 coding region in *I. scapularis* (Figure 2). We synthesized the upstream sequence incorporating the entire 5’ UTR up to the start codon and including the added restriction enzyme cloning sites, it resulted in a 1,301 bp fragment.

To test the promoter function, we transfected the HSP70 endogenous promoter construct with Renilla luciferase into pGL-Rluc into ISE6 cells, an *I. scapularis* embryonic cell line. The cells were lysed 9 days after transfection and analyzed for luciferase activity (Figure 3A). The initial results indicate that HSP70 is indeed active (Figure 3).

**Figure 3:**
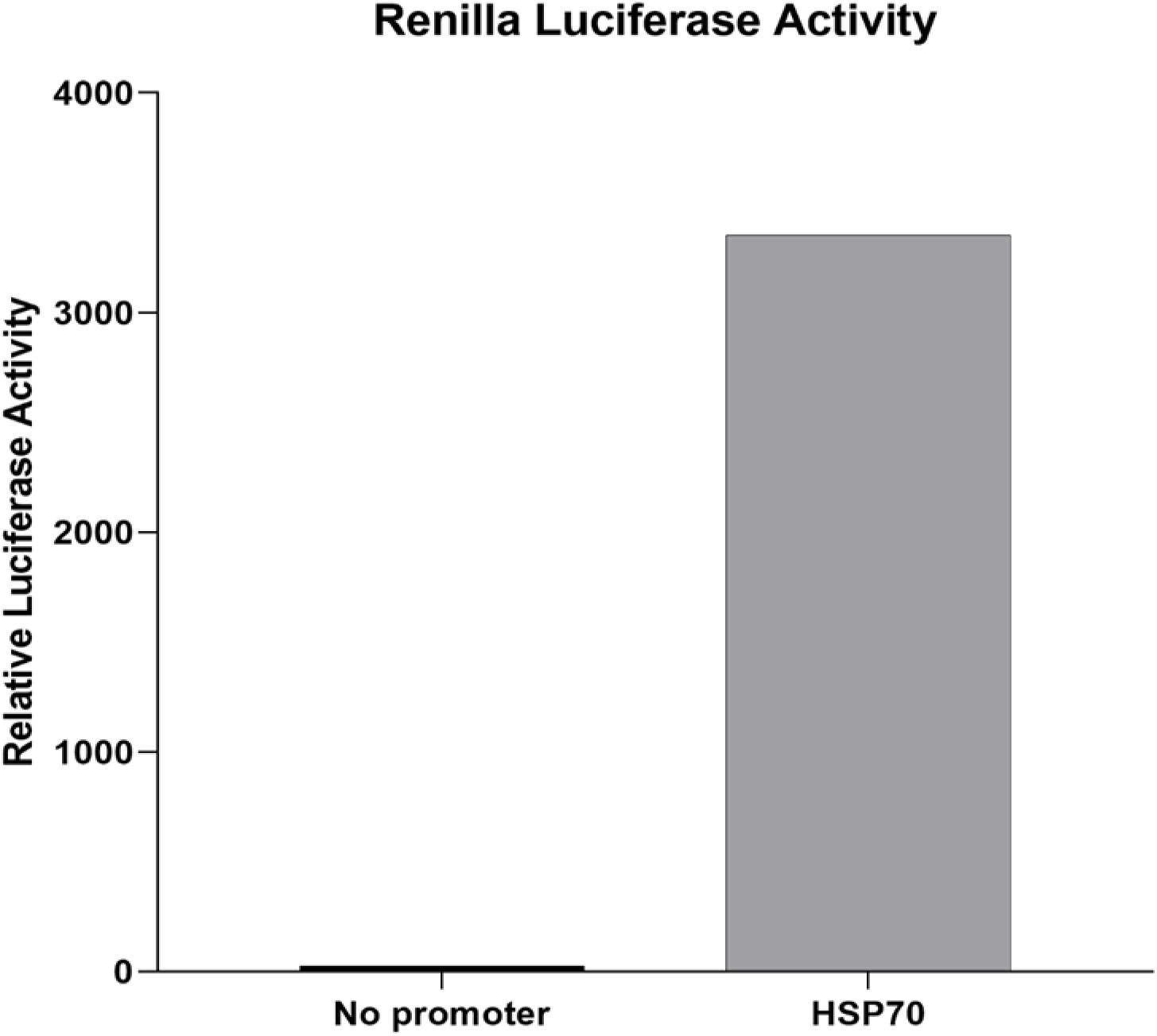
*Ixodes scapularis* ISE6 cell line transfected with various *Renilla* luciferase constructs. Cells were lysed, and lysates were examined for luciferase activity using a plate reader. Luminescence was measured at 10 s at Relative Luciferase Units.

### 3xP3 promoter

The 3xP3 promoter is a synthetic promoter that contains three Pax6 transcriptional activator homodimer binding sites. Multiple experiments with 3xP3-driven fluorescent constructs in ISE6 cells failed to show any fluorescence (data not shown), suggesting it is not functional in tick cells. The 3xP3 used in these prior experiments contained a *Drosophila* HSP70 minimal promoter (***Berghammer et al., 1999; Horn et al., 2000***). We swapped the minimal *Drosophila* HSP70 (123 bp) with part of the IsHSP70 (473 bp) from our confirmed functional construct (Figure 4A).

**Figure 4:**
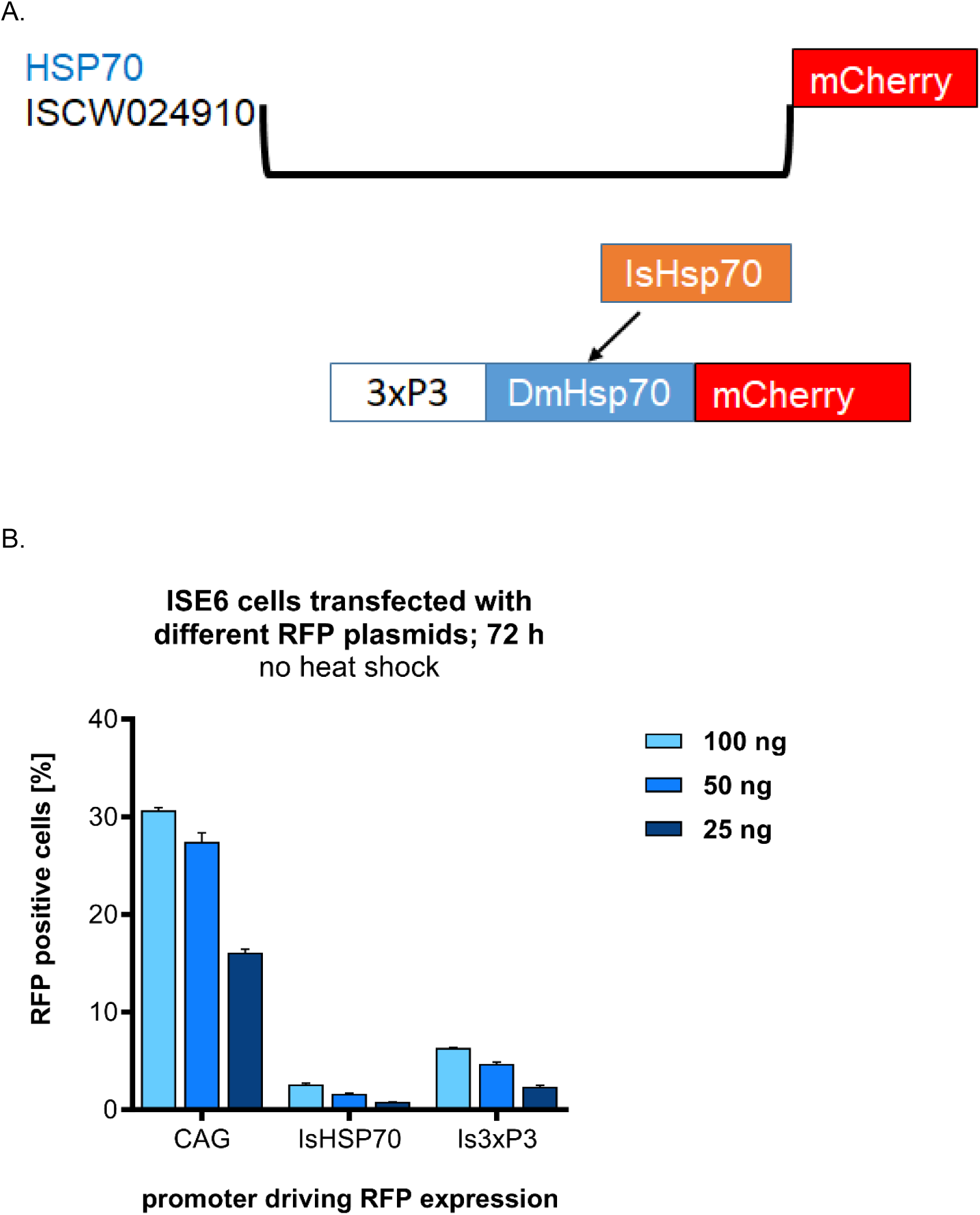
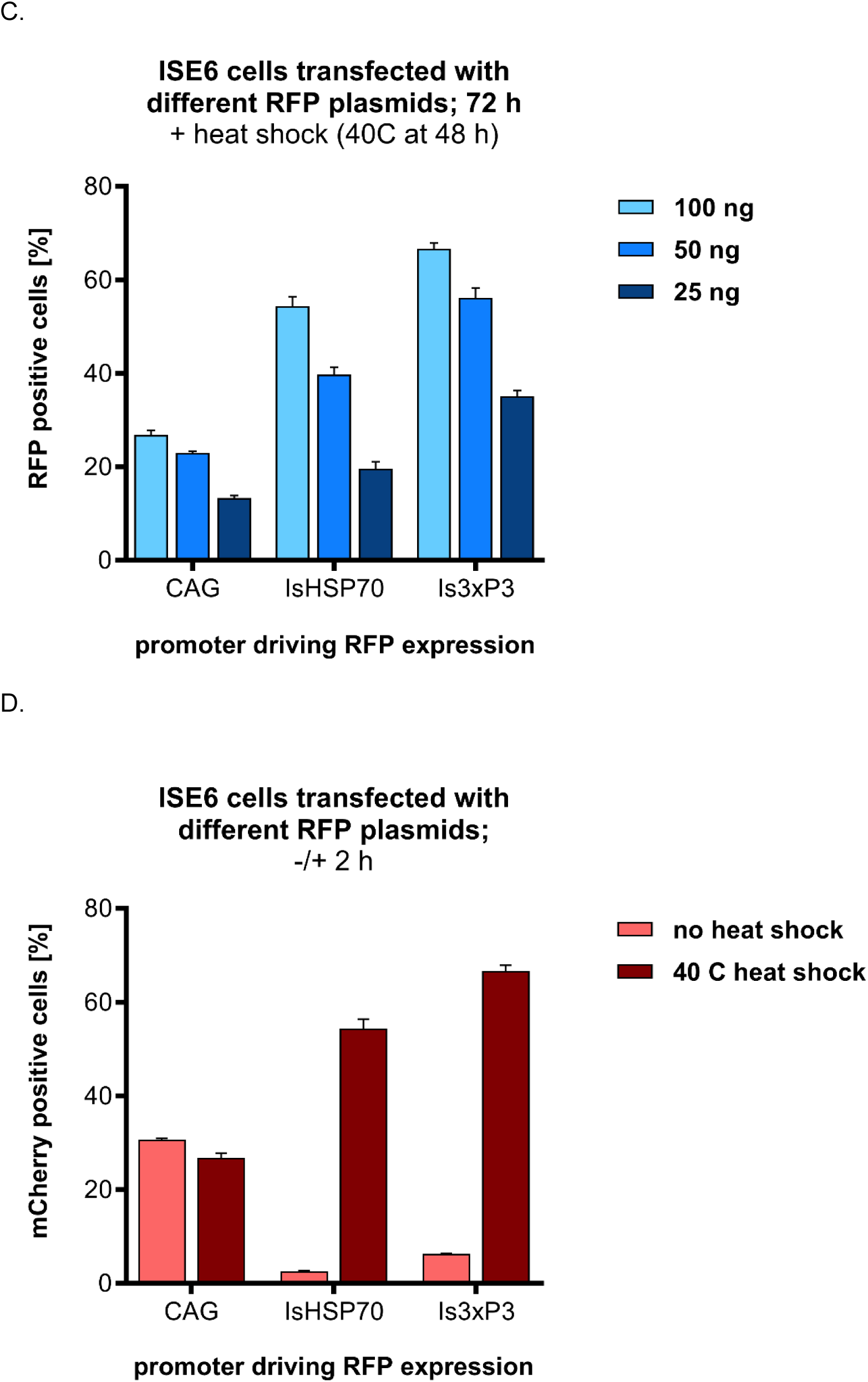
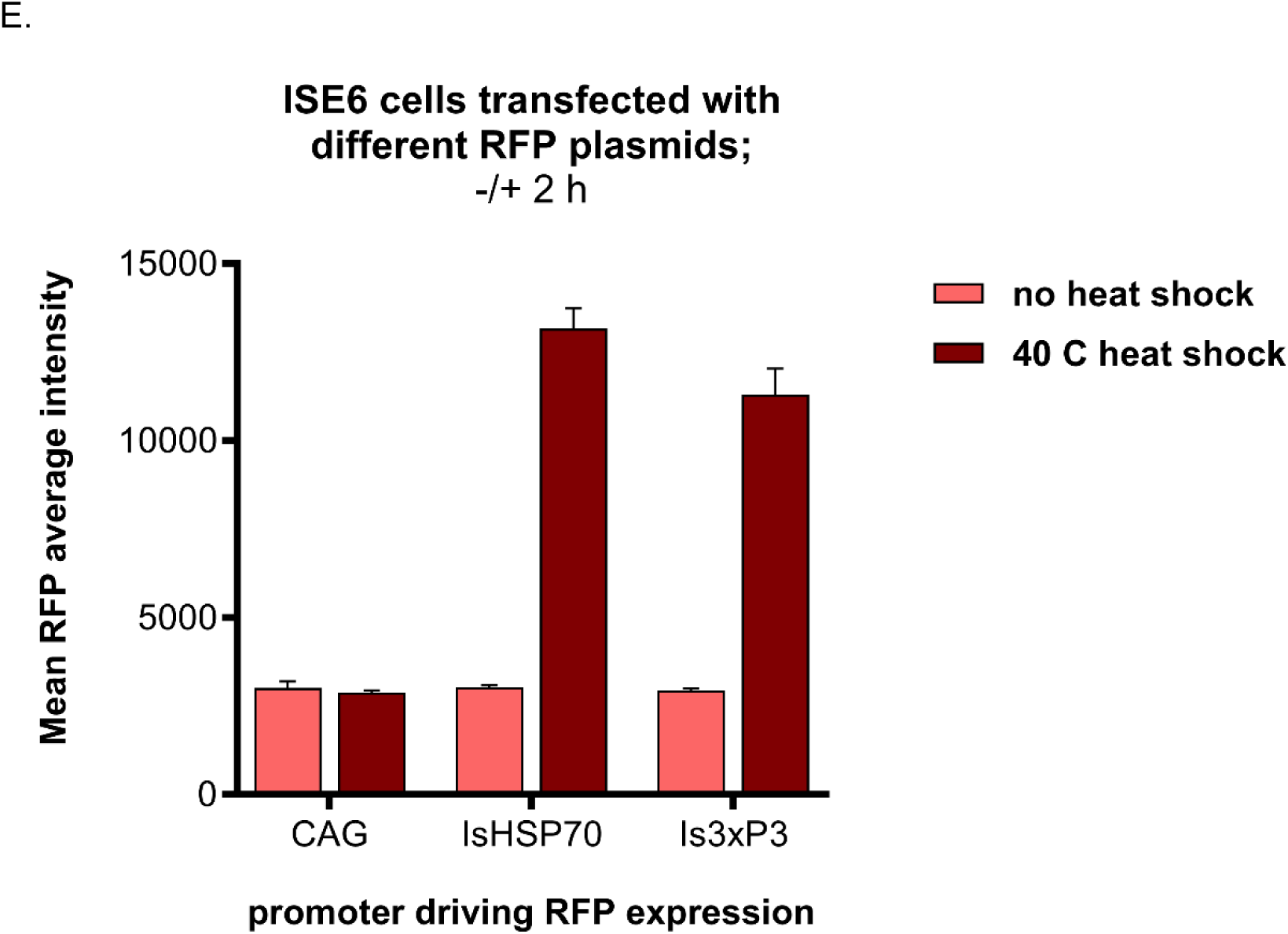
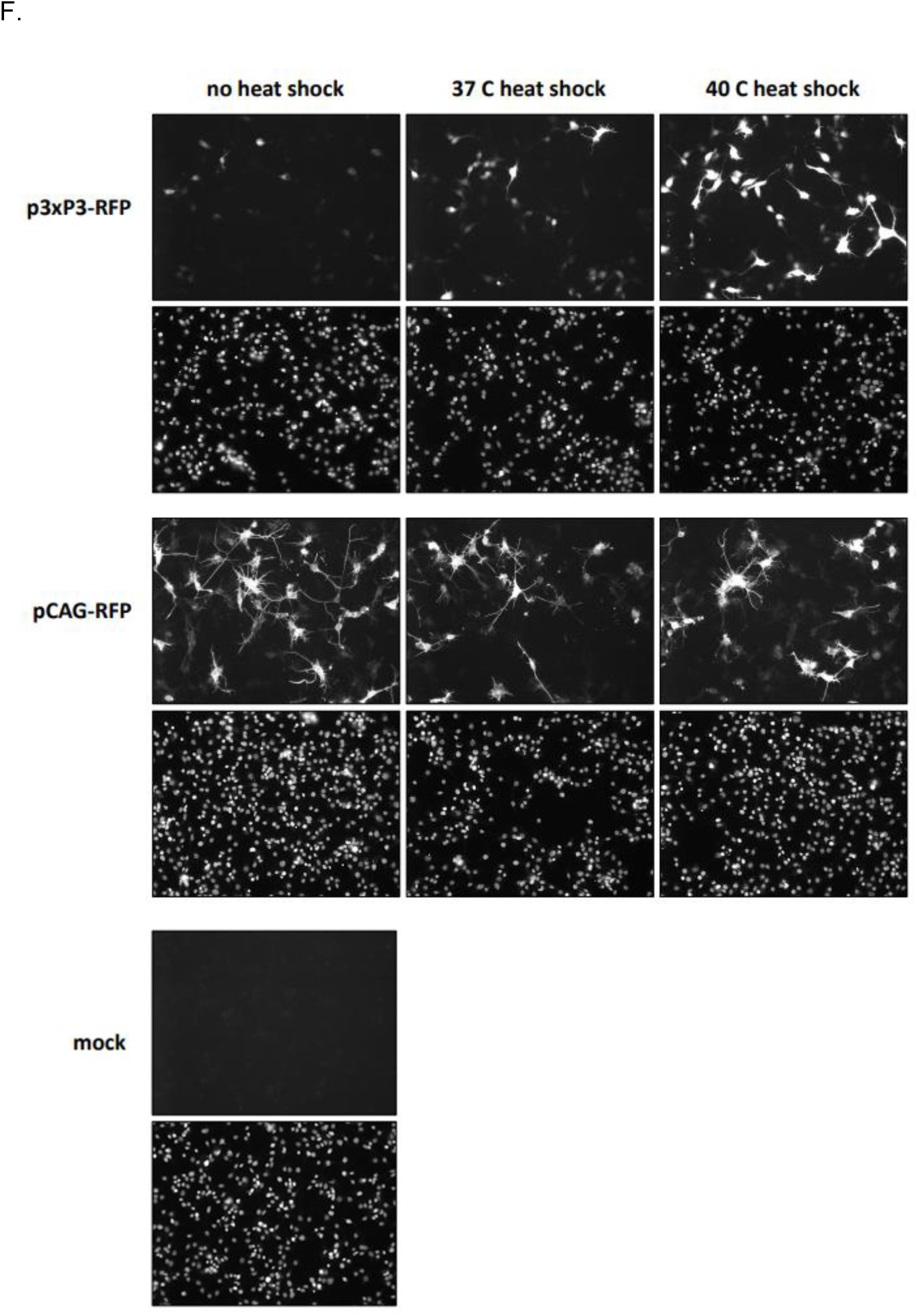
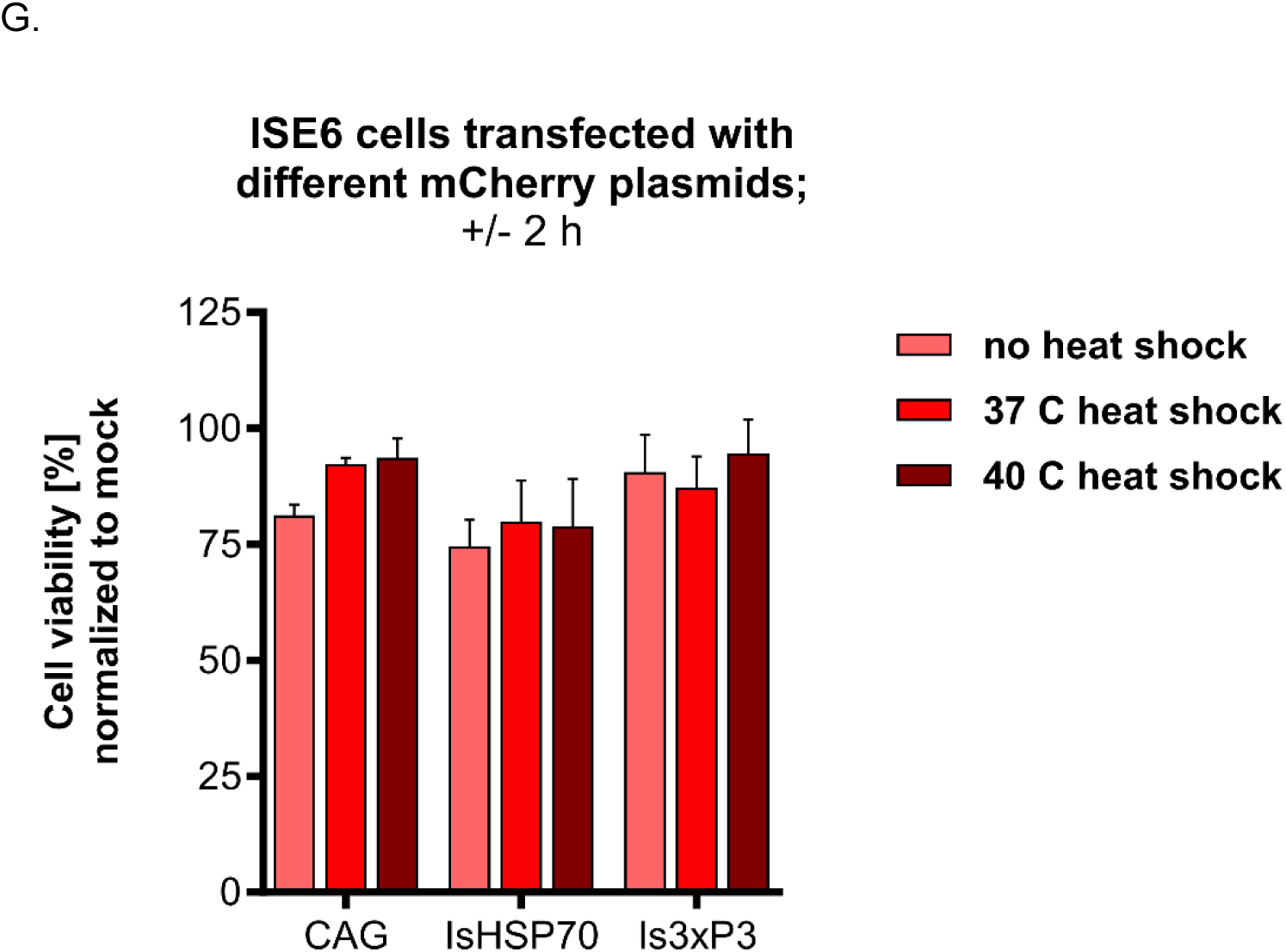
A: A gene map of swapped minimal *Drosophila* HSP70 (123 bp) with the *I. scapularis* HSP70 (473 bp). B: ISE6 cells transfected with 25, 50, or 100 ng of CAG-mCherry, IsHSP70-mcherry, and Is3xP3-mCherry constructs. Cells were fixed and imaged 72 h post transfection. C: ISE6 cells transfected with 25, 50, or 100 ng of CAG-mCherry, IsHSP70-mCherry, and Is3xP3-mCherry constructs. Transfected cells were subjected to a 2 h heat shock (40 °C) at 48 h post-transfection before being fixed 24 hours later (72 h post-transfection) and imaged. D: Percentage of fluorescent positive ISE6 cells transfected with 50 ng of CAG-, IsHSP70-, and Is3xP3-mCherry constructs. Cells were cultured at 32 °C (non-heat shock) or subjected for 2 h to 40 °C (heat shock) at 48 h post-transfection. Cells were then fixed and imaged 24 h later (72 h post transfection). E: Mean fluorescent intensity of ISE6 cells transfected in D. F: ISE6 cells were transfected with Is3xP3-mCherry and CAG-mCherry, subjected at 48 h post-transfection to a 2 h heat shock at either 37 °C or 40 °C and imaged 24 h later (72 h post-transfection). Top panel: images showing transfected cells, Bottom panel: cell nuclei stained with DAPI. Non-heat shock-treated cells transfected similarly were used as control. Cells treated with transfection reagents without the plasmids are shown as mock. G: Cell viability based on the number of nuclei of ISE6 cells transfected in F.

To test whether 3xP3 was functional, we transfected *I. scapularis* specific 3xP3 (Is3xP3)-mCherry, IsHSP70-mCherry, and CAG-mCherry (positive control) at increasing concentrations (25, 50, and 100 ng/well) into ISE6 cells. We observed fluorescence comparable to the HSP70 promoter, with approximately 5% fluorescent positive cells. However, the fluorescent signal was still lower compared to the CAG-mCherry construct (Figure 4B). We then conducted heat shock experiments with IsHSP70-mCherry and Is3xP3-mCherry by similarly transfecting ISE6 cells with increasing concentrations of plasmid DNA (25, 50, or 100 ng/well) followed by a 2 h heat shock at 40 after 48 h. While the control cells cultured continuously at 32 (non-heat shock conditions) were less than 5% fluorescent positive (Figure 4B), cells that were subjected to a heat shock showed an approximately 10-fold increase in fluorescent positive cells with both IsHSP70 and Is3xP3 (Figure 4C).

Using an intermediate concentration of plasmid DNA (50 ng/well), we tested if the changes in the percentage of fluorescent cells were accompanied by an increase in fluorescent intensity. An increase of approximately 10-fold in percent fluorescent positive cells (Figure 4D), which was accompanied by an approximately 4-fold increase in average fluorescence intensity was recorded (Figure 4E). Cell viability based on the number of cells/well was unaffected when comparing heat-shocked and non-heat-shocked cells both qualitatively (Figure 4F) and quantitatively (Figure 4G).

## Discussion

Here, we have shown two heat-inducible promoters functional in ISE6 cells: *I. scapularis* endogenous HSP70 (IsHSP70) and an *I. scapularis* specific artificial 3xP3 promoter (Is3xP3). Under non-heat shock conditions, IsHSP70 and Is3xP3 yield low percentages of fluorescent positive cells (∼5%). In contrast, under heat shock conditions such as 40 C, the percentage of fluorescent positive cells increases significantly by approximately 10-fold. This data suggests that nearly 10 times the basal percentage of cells are successfully transfected with both promoter constructs but do not sufficiently express mCherry protein to be detected. Our data also demonstrates that IsHSP70 and Is3xP3 have low basal activity under non-heat shock conditions, which readily increases during a heat shock response.

The artificial 3xP3 promoter, tested originally in *D. melanogaster*, contains three binding sites for Pax6/eyeless homodimers upstream to a TATA box. The Pax6 is an evolutionarily highly conserved system described as the master regulator of eye development throughout the animal kingdom (***Gehring, 2002***), which is consistent with the broad function of 3xP3 (***Berghammer et al., 1999; Sheng et al., 1997***) as a promoter for fluorescent protein genes. The 3xP3 promoter has been successfully used as an adult eye and ocelli marker for transgenesis in *Drosophila*, houseflies, beetles, butterflies, mosquitoes, as well as flatworms (***González-Estévez et al., 2003; Hediger et al., 2001; Horn et al., 2000; Kokoza et al., 2001; Lorenzen et al., 2003; Marcus et al., 2004***). Although the artificial 3xP3 promoter is widely used; it is not universal and is non-functional in organisms such as the tephritid fly (***Schetelig and Handler, 2013a***) and has potentially weaker functionality in horn flies (***Xu et al., 2016***). Similarly, our previous work showed that 3xP3 is non-functional in ticks. Therefore, we hypothesized that replacing the *Drosophila* HSP70 minimal promoter in 3xP3 with an endogenous tick IsHSP70 might overcome the functionality issues. Although *I. scapularis* is an eyeless tick, Pax6 (which binds to 3xP3) is expressed in ISE6 cells, *I. scapularis* ticks (***Gulia-Nuss et al., 2016***) as well as ticks such as *Rhipicephalus sanguines* (XM_037656705.1/XP_037512638) (***Nuss et al., 2021***), *H. longicornis*, and *Dermacentor silvarum* (XM_037709500I/XP_037565428) with fully formed eyes (***Jia et al., 2020***).

In addition to eyes, 3xP3 has been shown to promote expression in the larval nervous system, which has been highly useful in identifying silk moth transformants (***Thomas et al., 2002***). In mosquito *A. aegypti*, the 3xP3 promoter expresses in neural tissues and anal glands/digestive tract in addition to eyes (***Kokoza et al., 2001; Shin et al., 2003***). In one *D. suzukii* transgenic line, 3xP3 is expressed in the abdominal region (***Schetelig and Handler, 2013b***). In the marine crustacean, *Parhyale hawaiensis,* expression is not in the eyes but in cells at the posterior of the brain (***Goldstein and Srivastava, 2022; Pavlopoulos and Averof, 2005***). Therefore, Is3xP3 could drive the expression of transgenes in other tissues such as neural or digestive tract tissue. The experiments to test Is3xP3 expression in tick tissues are underway. Heat-inducible promoters are useful in controlling gene expression with applications including transgenesis and genome editing by allowing researchers to have temporal control of transposable elements and Cas9 in CRISPR expression systems. We expect the potential localized tissue expression of Is3xP3, similar to insect-based 3xP3, would be helpful for transgenic tick applications.

## Acknowledgments

This project was funded through NIH-NIAID grants R21AI128393, R212200536 to MG-N, R012200185 to MG-N and MP, R21AI153927 to H-HH, and Plymouth Hill Foundation, NY to MG-N.

